# Cortical network underlying speech production during delayed auditory feedback

**DOI:** 10.1101/2020.11.11.378471

**Authors:** Muge Ozker, Werner Doyle, Orrin Devinsky, Adeen Flinker

**Author notes:** **Corresponding Author:** Muge Ozker, **Email:**.

## Abstract

Hearing one’s own voice is critical for fluent speech production as it allows for the detection and correction of vocalization errors in real-time. This behavior known as the auditory feedback control of speech is impaired in various neurological disorders ranging from stuttering to aphasia, however the underlying neural mechanisms are still poorly understood. Computational models of speech motor control suggest that, during speech production, the brain uses an efference copy of the motor command to generate an internal estimate of the speech output. When actual feedback differs from this internal estimate, an error signal is generated to correct the estimate and subsequent motor commands to produce intended speech. We were able to localize these neural markers using electrocorticographic recordings from neurosurgical subjects during a delayed auditory feedback (DAF) paradigm. In this task, subjects hear their voice with a time delay as they produced words and sentences (similar to an echo on a conference call), which is well known to disrupt fluency by causing slow and stutter-like speech in humans. We observed a significant response enhancement in auditory cortex that scaled with the duration of feedback delay indicating an auditory speech error signal. Immediately following auditory cortex, dorsal precentral gyrus (dPreCG), a region that has not been implicated in auditory feedback processing before, exhibited a markedly similar response enhancement suggesting a tight coupling between the two regions. Critically, response enhancement in dPreCG occurred only when subjects profoundly slowed down their speech during articulation of long utterances due to a continuous mismatch between produced speech and reafferent feedback. These results suggest that dPreCG plays an essential role in updating the internal speech estimates to maintain fluency as well as coordinating the efference copy and auditory error signals during speech production.

## Introduction

Human speech production is strongly influenced by the auditory feedback it generates. When we speak, we continuously monitor our vocal output and adjust our vocalization to maintain fluency. For example, speakers involuntarily raise their voice to be more audible when auditory feedback is masked in the presence of background noise [1, 2]. Similarly, when speakers hear themselves with a delay (e.g. voice delays or echoes in teleconferencing), they compensate for the auditory feedback delay by slowing down and resetting their speech. This compensatory adjustment of human vocalization provides evidence for a mechanism which detects and corrects vocal errors in real-time. Abnormal auditory feedback control has been implicated in various disorders including stuttering, aphasia, Parkinson’s disease, autism spectrum disorder and schizophrenia [3–7], however the neural underpinnings of this dynamic system remain poorly understood.

Predictive models of speech motor control suggest that the brain generates an internal estimate of the speech output during speech production using an efference copy of the vocal-motor command. When there is a mismatch between this internal estimate and the perceived (reafferent) auditory feedback, the auditory response is enhanced to encode the mismatch. This auditory error signal is then relayed to vocal-motor regions for the real-time correction of vocalization in order to produce the intended speech [8–10].

In support of these models, electrophysiological studies in non-human primates demonstrated increased activity in auditory neurons when the frequency of the auditory feedback is shifted during vocalization [11]. Behavioral evidence in human studies showed that when formant frequencies of a vowel or the fundamental frequency (pitch) is shifted, speakers change their vocal output in the opposite direction of the shift to compensate for the spectral perturbation [12–14]. In line with non-human primate studies, human neurosurgical recordings as well as neuroimaging studies demonstrated that these feedback-induced vocal adjustments are accompanied by enhanced neural responses in auditory regions [15–17].

An alternative method to manipulating the spectral features of auditory feedback is altering its temporal features by delaying the voice onset in real time, termed “delayed auditory feedback (DAF)”. First described in the 1950s, DAF strongly disrupts speech fluency leading to slower speech rate, pauses, syllable repetitions and increased voice pitch or intensity [18–20]. Further, higher susceptibility to DAF occurs in autism spectrum disorder, non-fluent primary progressive aphasia, schizophrenia and other neurological disorders [4–6]. Interestingly, DAF improves speech fluency in individuals who stutter and is a therapeutic approach in speech therapy for stuttering and Parkinson’s Disease [21–23]. Although these behavioral effects have been widely studied in both normal and clinical groups, only a few neuroimaging studies have investigated the neural responses. Studies have demonstrated enhanced responses in bilateral posterior superior cortices during delayed feedback compared with normal auditory feedback conditions [24–26]. However, the exact temporal dynamics and spatial distribution of the cortical networks underlying speech production and reafferent feedback processing remain unknown.

To address this issue, we leveraged the excellent spatial and temporal resolution of electrocorticography (ECoG). Using ECoG, we acquired direct cortical recordings from 15 epilepsy patients while they read aloud words and sentences. As they spoke, we recorded their voice and played it back to them through earphones either simultaneously or with a delay (50, 100 and 200 milliseconds). Behaviorally, we found that subjects slowed down their speech to compensate for the delay, and did so more profoundly when producing sentences. Neurally, there was a significant increase in activity across a large speech network encompassing temporal, parietal and frontal sites that scaled with the duration of feedback delay. Critically, when speech was slowed down and became effortful, the dorsal division of the precentral gyrus was preferentially recruited at an early timing to support ongoing articulation. To our knowledge, we introduce the first temporally delayed feedback processing investigation with invasive human electrophysiology in which we reveal the fine-grained spatiotemporal dynamics of the neural mechanisms underlying compensatory adjustment of human vocalization.

## Results

Subjects (N = 15) performed a word-reading task (single 3-syllable words) while their voice onset was delayed (no delay, 50, 100 and 200 ms) and played back to them through earphones in real time, a paradigm known as delayed auditory feedback (DAF). We first analyzed the voice recordings of subjects and measured the articulation duration at different amount of delays to establish the behavioral effect of DAF (**Fig 1A**). Articulation duration increased slightly with delay: average articulation duration across subjects was 0.698, 0.726, 0.737 and 0.749 milliseconds for no delay, 50, 100 and 200 milliseconds delay conditions respectively. While this increase was not significant (**Fig 1B, ANOVA:** F = 1.985 p = 0.165) we observed robust neural changes.

**Fig 1.**
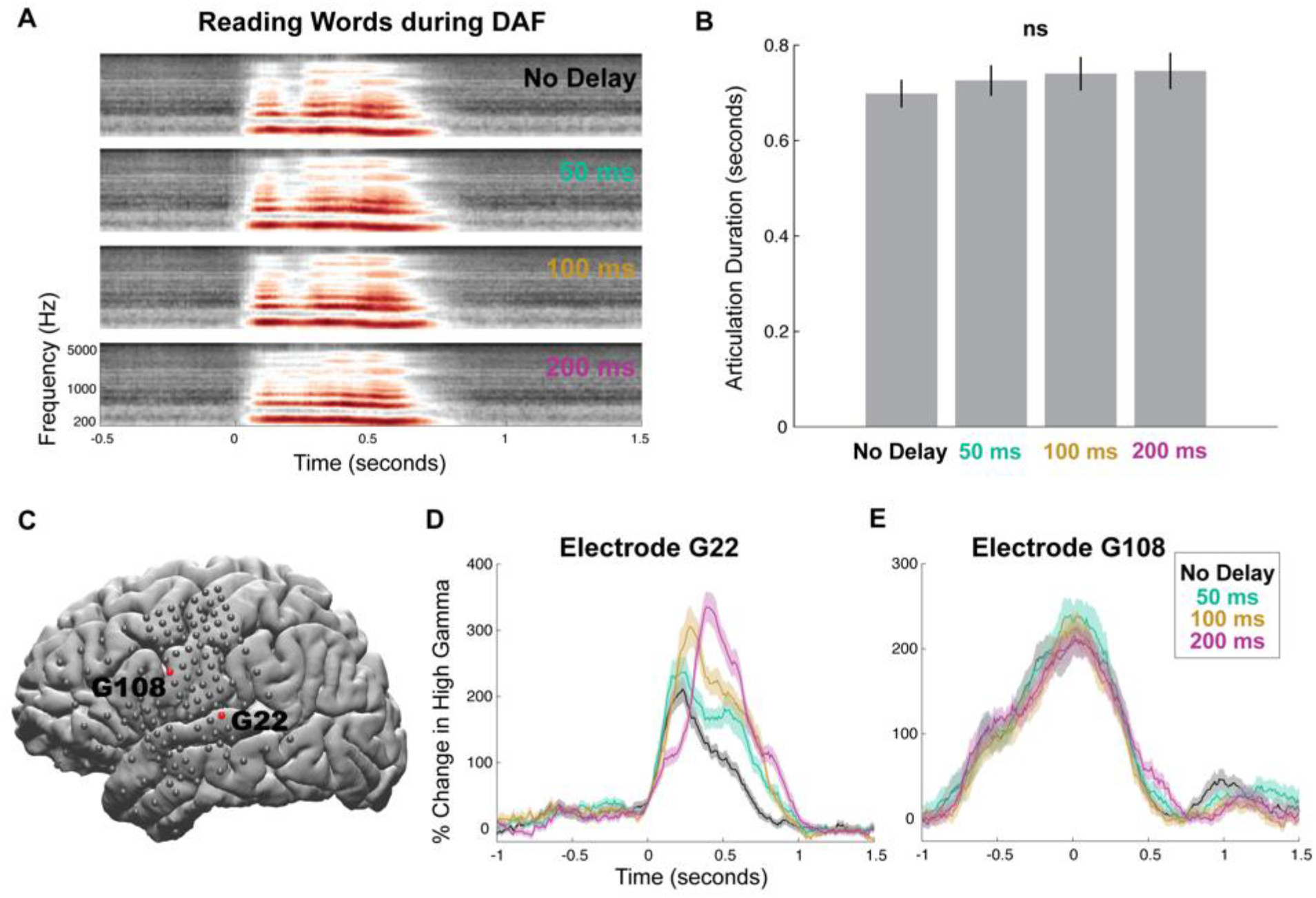
Behavioral and neural responses during word-reading with DAF. **(A)** Speech spectrogram of a single subject articulating words during DAF conditions. **(B)** Mean articulation duration of words during DAF conditions averaged across subjects. Error bars show SEM over subjects. **(C)** Cortical surface model of the left hemisphere brain of a single subject. Gray circles indicate the implanted electrodes. Red highlighted electrodes are located on the STG (G22) and on the vPreCG (G108). **(D)** High gamma responses in an auditory electrode (G22) to articulation of words during DAF conditions (color coded). Shaded regions indicate SEM over trials. **(E)** High gamma responses in a motor electrode (G108) to articulation of words during DAF conditions (color coded). Shaded regions indicate SEM over trials. **SEM** = Standard error of the mean

To quantify the neural response, we used the high gamma broadband signal (70-150 Hz, *see Methods*), a widely used index of cortical activity which correlates with underlying neuronal spike rates [30–32]. Two response patterns emerged among the electrodes that showed significant activity during speech production (*see Electrode Selection in Methods*). In the first pattern, shown on a representative auditory electrode located in the STG (**Fig 1C**), neural response started after speech onset and its amplitude increased significantly with delay (**Fig 1D, ANOVA:** F = 37, p = 1.55×10^-8^). In the second pattern, shown on a representative motor electrode located in vPreCG (**Fig 1C**), neural response started before speech onset and its amplitude was not affected by delay (**Fig 1E, ANOVA:** F = 0.084, p = 0.772). This result demonstrated that although DAF did not significantly affect speech behavior (i.e. articulation duration), it affected the neural response in auditory sites that are involved in speech processing.

To characterize the two major response patterns in the brain we chose to use an unbiased, data-driven approach which does not impose any assumptions or restrictions on the selection of responses. We performed an unsupervised clustering analysis using the NMF algorithm on neural responses across all delay conditions, brain sites and subjects [29, 33]. The clustering analysis identified the major response patterns represented by two distinct clusters, which corroborated our representative results shown in a single subject (**Fig 1C-E**) as well as visual inspection of the data across subjects. The first response pattern (Cluster 1, N = 125 electrodes) started after speech onset and peaked at 320 ms reaching 115 percent change in amplitude. The second response pattern (Cluster 2, N = 253 electrodes) started much earlier approximately 750 ms prior to speech onset and peaked 140 ms after speech onset reaching 60 percent change in amplitude (**Fig 2A**). These two clusters had a distinct anatomical distribution (**Fig 2B**): Cluster 1 was mainly localized to STG suggesting an auditory function while Cluster 2 was localized to frontal cortices suggesting a pre-motor and motor function.

**Fig 2.**
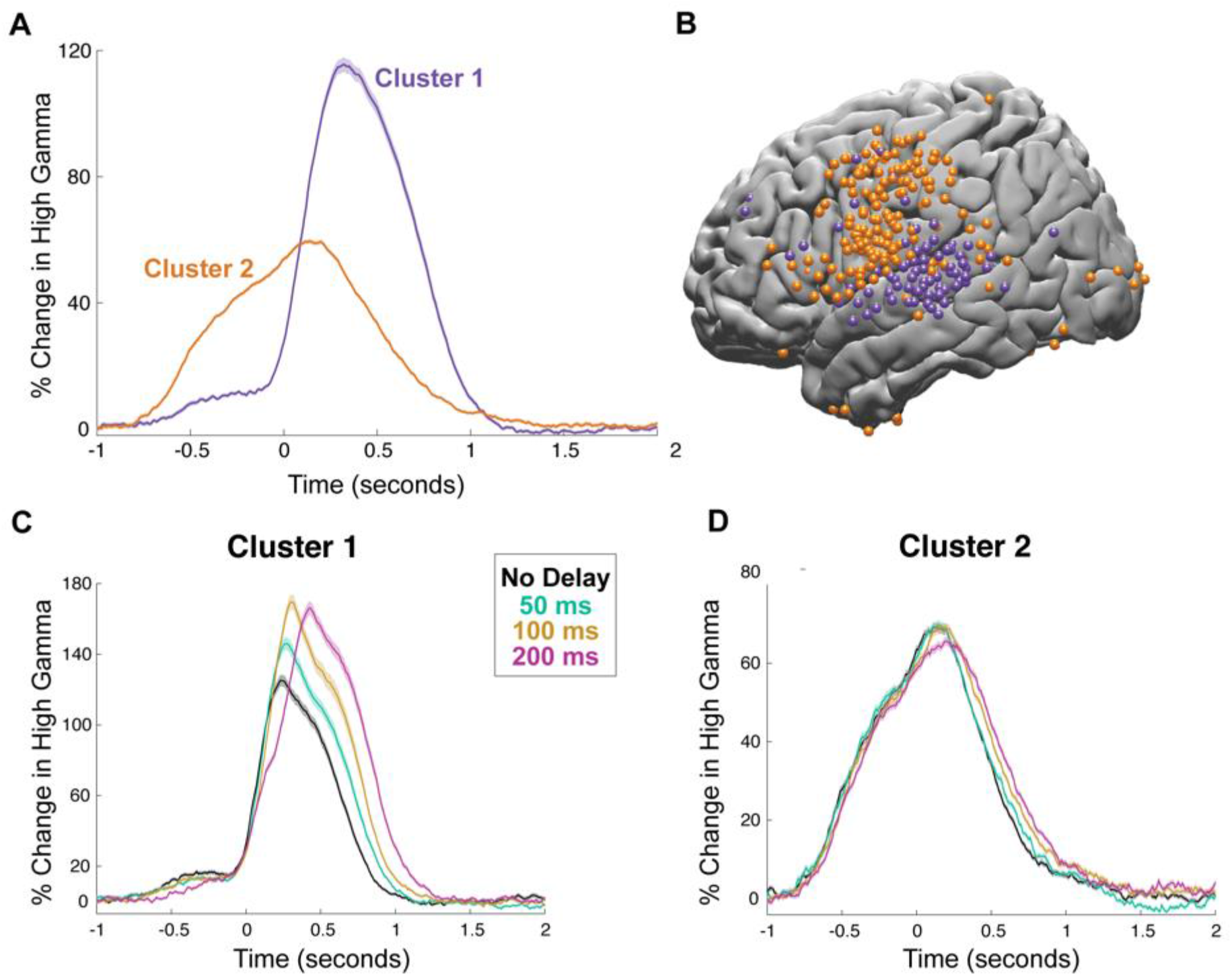
Clustering with non-negative matrix factorization. **(A)** High gamma responses averaged across electrodes in the two clusters provided by the unsupervised NMF. Shaded regions indicate SEM over trials. **(B)** Spatial distribution on cortex of electrodes in the two clusters displayed on the left hemisphere of a template brain. **(C)** High gamma responses to articulation of words during DAF conditions averaged across electrodes in Cluster 1. Shaded regions indicate SEM over trials. **(D)** High gamma response to articulation of words during DAF averaged across electrodes in Cluster 2. Shaded regions indicate SEM over trials. **SEM** = Standard error of the mean

Next, we examined the effect of DAF on these two clusters. The amplitude of the neural response increased significantly with delay in Cluster 1 (**Fig 2C, ANOVA:** F = 5.35, p = 0.02), but not in Cluster 2 (**Fig 2D, ANOVA:** F = 1.65, p = 0.2). The duration of the neural response did not show a significant increase in either of the clusters (**ANOVA:** F = 1, p = 0.32 for Cluster 1 and F = 0.01, p = 0.92 for Cluster 2).

Reading words with DAF did not prolong articulation duration and while it increased neural responses in auditory regions, it did not affect responses in motor regions. We hypothesized that a longer and more complex stimulus may elicit a stronger behavioral response and motor regions will show an effect of DAF when articulation is strongly affected. To test this prediction, we performed another experiment in which subjects read aloud sentences during DAF. Indeed, articulating longer speech segments (8-word sentences) during DAF resulted in a significantly stronger behavioral effect (**Fig 3A**). Articulation duration increased significantly with delay: average articulation duration across subjects was 2.761, 2.942, 3.214 and 3.418 seconds for no delay, 50, 100 and 200 milliseconds delay conditions respectively (**Fig 3B, ANOVA:** F = 17.11, p = 0.0001).

**Fig 3.**
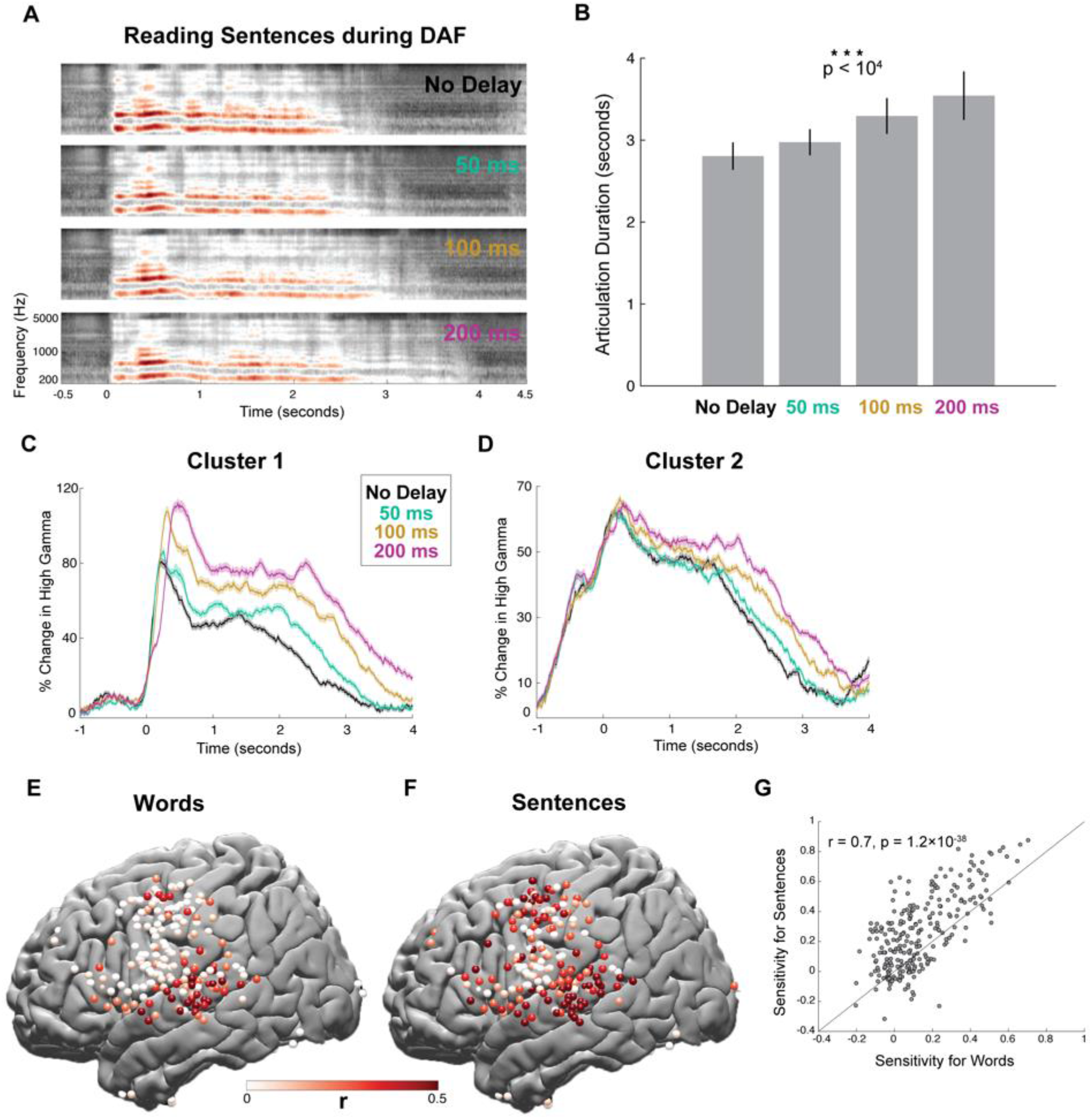
Behavioral and neural responses during sentence-reading with DAF. **(A)** Speech spectrogram of a single subject articulating sentences during DAF conditions showing a marked increase in articulation duration. **(B)** Mean articulation duration of sentences during DAF conditions averaged across subjects showing a significant effect of duration. Error bars show SEM over subjects. **(C)** High gamma responses to articulation of sentences during DAF conditions averaged across electrodes in Cluster 1. Shaded regions indicate SEM over trials. **(D)** High gamma responses to articulation of sentences during DAF conditions averaged across electrodes in Cluster 2. Shaded regions indicate SEM over trials. **(E)** Anatomical map of electrodes across all subjects displayed on the left hemisphere of a template brain showing the neural sensitivity to DAF during word-reading. **(F)** Anatomical map of electrodes across all subjects displayed on the left hemisphere of a template brain showing the neural sensitivity to DAF during sentences-reading. **(G)** Scatter plot and fitted regression showing significant correlation between sensitivity to DAF for the word-reading and sentence-reading tasks. Each circle represents one electrode. **SEM** = Standard error of the mean

Next, we examined the neural response to DAF in the two electrode clusters we identified previously (**Fig 2B**). When reading words during DAF, amplitude of the neural response increased with delay in Cluster 1 but not in Cluster 2 (**Fig 2C and 2D**). However, when reading sentences during DAF, neural response in both clusters showed a sustained effect (**Fig 3C and 3D, ANOVA:** F = 18, p = 2.95 ×10^-5^ for Cluster 1 and F = 4.8, p = 0.03 for Cluster 2). Also, when reading words during DAF, duration of the neural response in neither of the clusters showed a significant effect of delay. However, when reading sentences during DAF, neural response duration in both clusters increased significantly with delay paralleling the significant behavioral effect of DAF on articulation duration (**ANOVA:** F = 21.6, p = 10^-5^ for Cluster 1 and F = 35.5, p = 10^-8^ for Cluster 2).

Our clustering analysis identified two response components that were mostly anatomically distinct reflecting an auditory response to self-generated speech and a motor response to articulation. The auditory component was unique in exhibiting an enhanced response during both word-reading and sentence-reading with DAF likely representing an auditory error signal. Response enhancement changed as a function of feedback delay, which suggests that auditory error signal does not simply encode the mismatch between intended and perceived speech but is sensitive to the amount of mismatch. We quantified the error signal by calculating a sensitivity index for each electrode by measuring the trial-by-trial correlation between the delay and the neural response averaged over a 0 to 1 s time window for words and over 0 to 3 s for sentences. A large sensitivity value indicated a strong response enhancement with increasing delays.

Comparing sensitivity indices for word-reading and sentence-reading tasks revealed that electrodes that were sensitive to DAF for the word-reading task were also sensitive for the sentence-reading task (r = 0.7, p = 1.2×10^-38^, **Fig 3G**), however the majority of electrodes showed larger sensitivity to DAF for the sentence-reading task. Moreover, several sites such as IFG and dPreCD showed increased sensitivity in the sentence-reading task (**Fig 3E-F**). This result suggests that articulating longer speech segments during DAF results in stronger overall sensitivity across auditory and motor regions and engages a larger brain network uniquely recruiting additional frontal regions.

We further examined the neural response to DAF in six different regions of interest based on within subject anatomy: STG, vPreCG, dPreCG, postCG, SMG and IFG (**Fig 4A-F**). Comparing sensitivity indices for word-reading and sentence-reading tasks in these regions revealed that all six regions showed larger sensitivity to DAF during sentence-reading (unpaired ttest; STG: t = 3.13, p = 0.0025, vPreCG: t = 4.55, p = 2.7×10^-5^, dPreCG: t = 5.15, p = 3×10^-6^, postCG: t = 3.1, p = 0.0024, SMG: t = 1.87, p = 0.07, IFG: t = 2.89, p = 0.0069; **Fig 4G**). To reveal how response enhancement to DAF changed across time during the sentence-reading task, we performed a one-way ANOVA at each time point (*see Methods*) and marked the timepoints when the neural response to the four delay conditions were significantly different for at least 200 consecutive milliseconds. Significant divergence onset during sentence-reading started the earliest in STG at 80 ms after speech onset, followed by dPreCG at 350 ms and SMG gyrus at 680 ms, and lasted throughout the stimulus. In postCG, vPreCG and IFG, responses diverged much later, at 1.82, 1.87 and 2.30 s respectively (**Fig 4H**). These timings reflect when cortical regions become sensitive to DAF and provide evidence for two distinct timeframes of early (STG, dPreCG, SMG) and late (postCG, vPreCG, IFG) recruitment.

**Fig 4.**
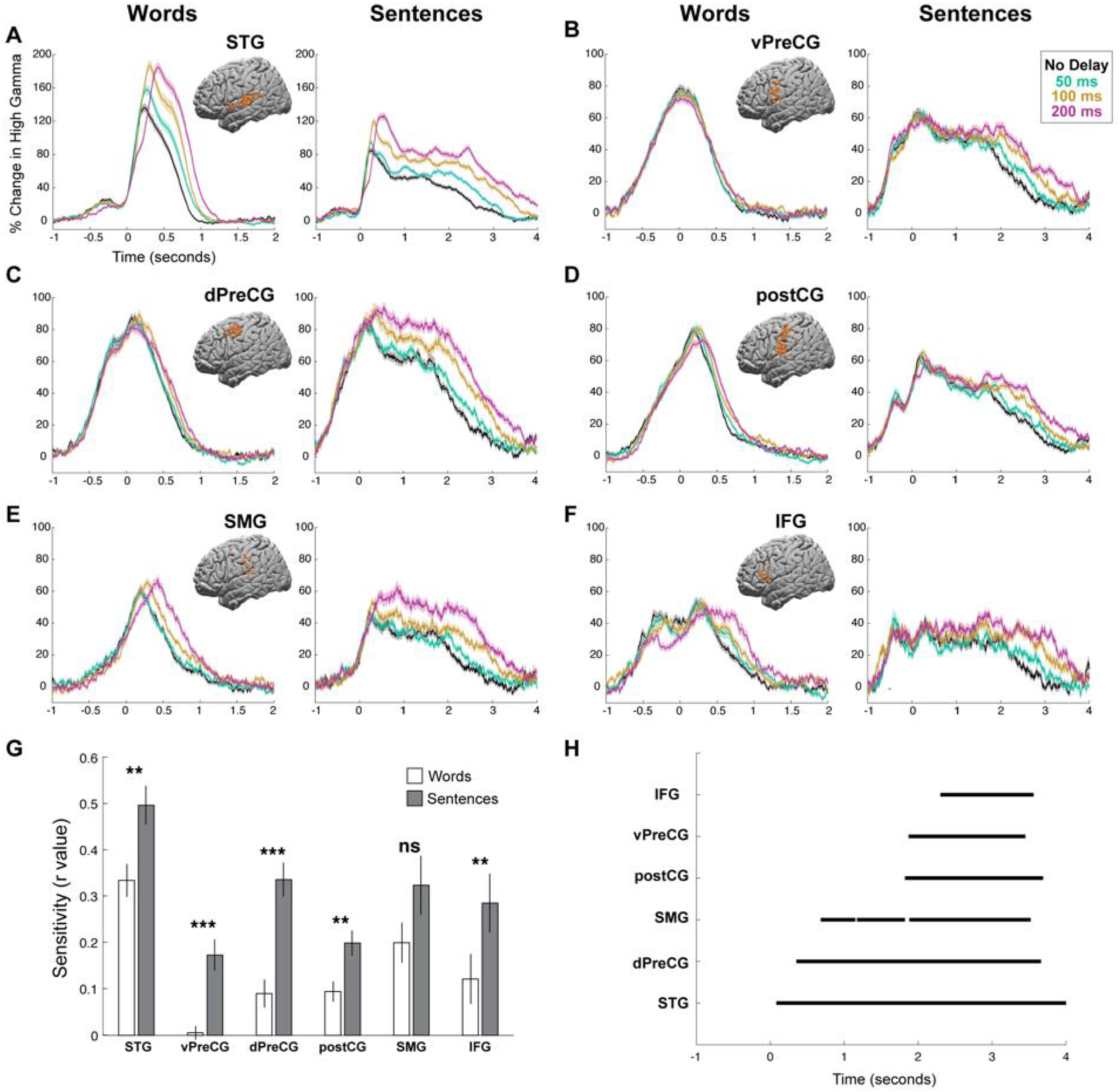
Neural responses to DAF by regions. **(A-F)** High gamma responses to articulation of words and sentences during DAF in six different regions: STG (a), vPreCG (b), dPreCG (c), postCG (d), SMG (e) and IFG (f). Inset brain figures shows the location of electrodes across all subjects on the left hemisphere of a template brain. Colors represent the various DAF conditions and shaded regions indicate SEM over trials. **(G)** Sensitivity to DAF during word-reading and sentence-reading tasks averaged across electrodes (error bars indicate SEM over electrodes) in six different regions. **(H)** Time intervals when the neural response to reading sentences during DAF diverged significantly across conditions within each of the six different regions. Significance is assessed using a shuffled permutation approach. **SEM** = Standard error of the mean

Examining different regions of the speech network revealed variable degree of neural response enhancement to DAF. The increase in response amplitude was usually accompanied by an increase in response duration, which was a result of longer articulation duration. In order to disentangle the enhanced neural response representing an error signal with longer articulation duration due to the exerted behavior, we applied a temporal normalization technique. We transformed the neural response time series using dynamic time warping (DTW) so that they would match in time span. DTW measures the similarity between two temporal sequences with different lengths by estimating a distance metric (a warping path) that would transform and align them in time. Matching the neural responses in time allowed us to directly compare their amplitudes and identify which brain regions produce an error signal in response to DAF rather than just sustained activity in time due to longer articulation.

We compared two conditions, which show the largest neural response difference in terms of amplitude and duration; 0 and 200 ms delay conditions (see *Dynamic Time Warping Analysis in Methods*). After aligning the responses in time, we averaged the amplitudes of the time-warped signals over time (0-6 s) and compared the two conditions by running an unpaired t-test. Amplitudes of the time-warped responses to DAF were significantly larger in STG (t = 5.5, p = 7.7×10^-8^), SMG (t = 3.06, p = 0.0025), dPreCG (t = 2.4, p = 0.016) and IFG (t = 3.4, p = 10^-3^) but not in vPreCG (t = 0.82, p = 0.41) and postCG (t = 1.66, p = 0.0971) regions (**Fig 5A-F**). Lastly, to examine the spatial distribution of the (time-warped) error response in more detail, we calculated the percent increase in response amplitude in single electrodes (HGB_200_ - HGB_no delay_ / HGB_no delay_*100). This analysis revealed that the magnitude of the error response was variable both across and within the regions of the speech network. Overall the error signal centered around four major cortical networks: STG, IFG, SMG and dPreCG (**Fig 5G**).

**Fig 5.**
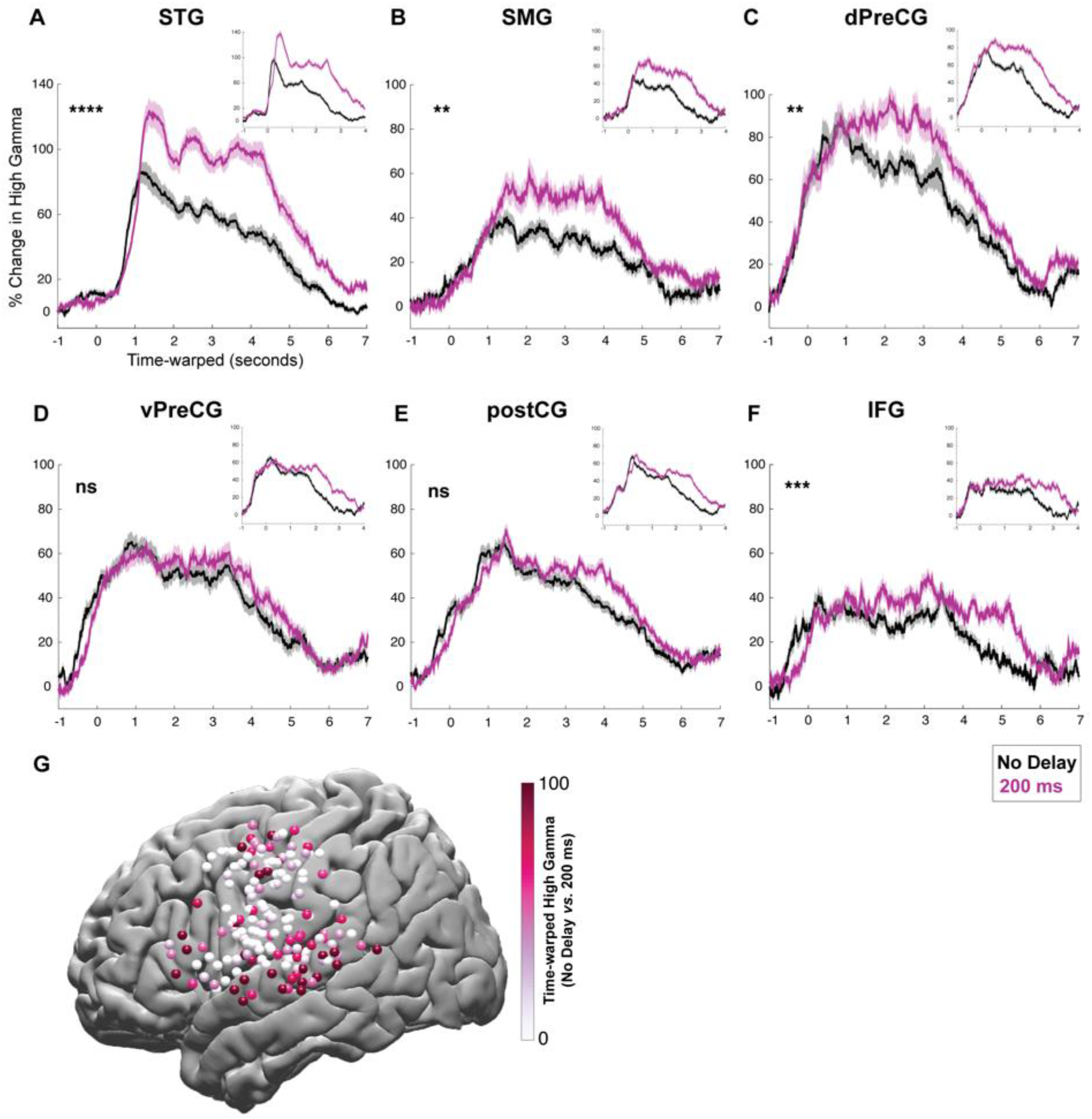
Time warped neural responses during sentence-reading with DAF. **(A-F)** High gamma responses after correction for articulation duration using dynamic time warping. Activity locked to articulation of sentences is shown for no delay (black) and 200 ms delay (magenta) conditions in six different regions: STG, SMG, vPreCG, dPreCG, postCG and IFG. Inset figures show the uncorrected high gamma responses which include the normal articulation timing. **(G)** Spatial distribution of the increase in neural responses to sentence-reading during DAF (200 ms) compared to no delay condition. Electrodes across all subjects are displayed on the left hemisphere of a template brain.

## Discussion

Artificially slowing down speech when hearing one’s own delayed voice provides a strong framework to investigate how auditory feedback influences the motor control of speech. Our study is one of the few electrophysiological investigations [34, 35], and to our knowledge, the only ECoG investigation of the underlying neural substrates. We compared the effects of DAF on producing short versus long speech segments by using word and sentence stimuli and showed that producing sentences during DAF had a stronger disruptive effect on speech. We used an unsupervised clustering algorithm (NMF) to determine auditory and motor regions involved in speech production and then identified four subregions of the speech network that are centrally engaged in the processing of auditory feedback: STG, SMG, dPreCG and IFG. The exquisite resolution of ECoG provided us with the precise spatiotemporal evolution of feedback processing in these distinct regions. Neural responses were enhanced in amplitude and extended in duration for large delays reflecting the error signal caused by altered feedback and the subsequent longer articulation. To dissociate the error signal from the effect of prolonged articulation, we used dynamic time warping algorithm and temporally aligned the neural signals with the patients’ speech acoustics. We found that dPreCG showed response enhancement immediately after auditory cortex when speech fluency was strongly disrupted during production of sentences with DAF. These results highlighted dPreCG as a critical region for maintaining speech fluency when dynamic auditory feedback processing is required to produce longer utterances.

During speech production, the reafferent (perceived) auditory feedback is not immediately useful due to noise and delays in neural processing (axonal transmission, synaptic processes etc.). According to predictive models of speech motor control, the brain must therefore rely on an internal estimate of the auditory feedback and use reafferent feedback only to correct this internal estimate. When an utterance is produced, an efference copy of the motor command is used to make a prediction of the current articulatory state (e.g. state of the vocal tract) and the subsequent sensory outcome. As long as there is no mismatch between the predicted and the reafferent feedback, the brain can rely on its internal estimate. However, when there is a mismatch, the brain generates an error signal to correct its internal estimate and the necessary motor commands to produce the intended utterance. In our paradigm, an artificially introduced delay generates a continuous mismatch between the predicted and the reafferent feedback. Specifically, the reafferent feedback matches the previous articulatory state rather than the current articulatory state. In this case, the issuing of new motor commands must be delayed, which would explain the slowing down of speech production. We found that producing single words with DAF elicited a nominal slowing down effect and increased neural responses only in auditory but not in motor regions. However, when subjects produced sentences with DAF, this longer and more complex stimulus elicited a much prominent slowing down effect and increased neural responses in auditory as well as motor regions.

Neural responses to DAF were enhanced in STG, SMG, dPreCG and IFG, which are anatomically connected by one of the major language pathways known as the superior longitudinal fasciculus [36, 37]. These regions are typically modeled as the main components of the dorsal stream for speech that is responsible for sensorimotor integration and auditory feedback processing [38]. In support of these theoretical models, clinical reports demonstrated that posterior STG and SMG damage are implicated in conduction aphasia [39] and patients with conduction aphasia are less affected by DAF [40] indicating the involvement of this regions in feedback processing.

Our analysis of the time course of responses revealed that response enhancement to DAF started in STG and closely followed by dPreCG providing further evidence for a functional correspondence between the two regions. dPreCG is a complex functional region implicated in auditory, motor and visual speech processing [41–44]. It is known to be activated not only during speaking but also during passive listening suggesting a role in mapping acoustic speech features onto the corresponding articulatory movements. We predict that dPreCG may be the hub for generating the internal estimate of the speech output by processing both the efference copy and auditory error signals during speech production.

Interestingly, response enhancement in dPreCG during auditory feedback processing has never been reported previously. Studies that altered auditory feedback using pitch perturbation demonstrated enhanced responses in ventral parts of the precentral gyrus, which correlated with the compensatory vocal adjustments [12–14]. In our experiments, vPreCG did not show any response enhancement neither for word nor for sentence production with DAF. It could be the case that vocal control that requires pitch adjustments activate vPreCG, while vocal control that requires long-term maintenance of other prosodic features such as tempo, rhythm and pause, activate dPreCG. dPreCG is well known to be involved in movement planning and execution [45, 46]. Previous studies reported increased activation in this region when subjects produced complex syllable sequences, suggesting a role in the planning and production of long speech utterances with appropriate syllable timing [47]. Considering that dPreCG activation occurred only when subjects produced sentences with DAF, we predict that it plays a critical role in maintaining prosody and speech fluency during the articulation of long utterances.

Our results showed that the maximal disruption of speech occurred at 200 ms feedback delay, which agrees with previous behavioral reports [18–20, 48, 49]. The dynamics of the cortical speech network can provide an explanation for this effect. Previous ECoG studies showed that IFG is activated before articulation onset and remained silent during articulation, while motor cortex is activated both before and during articulation. These studies suggested that IFG produces an articulatory code that is subsequently implemented by the motor cortex and reported a ~200 ms temporal lag between IFG and motor cortex activation [50, 51]. A feedback delay in the same order of this temporal lag likely interrupts propagation of the articulatory code from IFG to motor cortex, thereby disrupting speech. Unlike prior reports, we found sustained IFG activity throughout speech production, however this was during DAF where sustained IFG recruitment may be necessary to support compensatory speech correction. The onset of IFG activation was seen in conjunction with dPreCG, however sensitivity to DAF was seen at two distinct time periods with an early recruitment of dPreCG and much later involvement of IFG.

Behavioral paradigms that manipulate auditory feedback have been widely used for decades to understand speech motor control, however the cortical dynamics underlying this process remained largely unknown. We elucidate the magnitude, timing and spatial distribution of the neural responses that encode the mismatch between produced speech and its perceived feedback. Our results highlighted STG, SMG, IFG and dPreCG as critical players in detection and correction of vocalization errors. Specifically, we find that dPreCG is a selective region which is recruited immediately when auditory feedback becomes unreliable and production more effortful, implicating it in auditory-motor mapping that underlies vocal monitoring of human speech.

## Materials and methods

### Subject information

All experimental procedures were approved by the NYU Grossman School of Medicine Institutional Review Board. The review board considered: 1) the risks and anticipated benefits (if any) to subjects 2) the selection of subjects 3) the procedures for securing and documenting informed consent 4) the safety of subjects 5) the privacy of subjects and confidentiality of the data. The approval is for study i14-02101 and NYU Grossman School of Medicine Federal wide Assurance number is FWA00004952. All patients provided written consent prior to the study as well as an additional oral consent at the start of the experiment.

15 neurosurgical epilepsy patients (8 females, mean age: 34, 2 right, 9 left and 4 bilateral hemisphere coverage) implanted with subdural and depth electrodes provided informed consent to participate in the research protocol. Electrode implantation and location were guided solely by clinical requirements. 3 patients were consented separately for higher density clinical grid implantation, which provided denser sampling of underlying cortex.

### Experiment setup

Subjects were tested while resting in their hospital bed in the epilepsy-monitoring unit. Visual stimuli were presented on a laptop screen positioned at a comfortable distance from the subject. Auditory stimuli were presented through earphones (Bed Phones On-Ear Sleep Headphones Generation 3) and subjects’ voice was recorded using an external microphone (Zoom H1 Handy Recorder).

### Delayed auditory feedback experiment

The experiment consisted of a word-reading session and a sentence-reading session. 10 different 3-syllable words (e.g. document) were used in the word-reading session, and 6 different 8-word sentences (e.g. The cereal was fortified with vitamins and nutrients) were used in the sentence-reading session. Text stimuli were visually presented on the screen and subjects were instructed to read them out loud. As subjects spoke, their voices were recorded using the laptop’s internal microphone, delayed at 4 different amounts (no delay, 50, 100, 200ms) using custom script (MATLAB, Psychtoolbox-3) and played back to them through earphones. A TTL pulse marking the onset of a stimulus, the delayed feedback voice signal (what the patient heard) and the actual microphone signal (what the patient spoke) were fed in to the EEG amplifier as an auxiliary input in order to acquire them in sync with the EEG samples. Trials, which consisted of different stimulus-delay combinations, were presented randomly (3 to 8 repetitions) with a 1 second inter-trial-interval.

### Electrocorticography (ECoG) recording

ECoG was recorded from implanted subdural platinum-iridium electrodes embedded in flexible silicon sheets (2.3 mm diameter exposed surface, 8 x 8 grid arrays and 4 to 12 contact linear strips, 10 mm center-to-center spacing, Ad-Tech Medical Instrument, Racine, WI) and penetrating depth electrodes (1.1 mm diameter, 5-10 mm center-to-center spacing 1 x 8 or 1 x 12 contacts, Ad-Tech Medical Instrument, Racine, WI). Three subjects consented to a research hybrid grid implanted which included 64 additional electrodes between the standard clinical contacts (16 × 8 grid with sixty-four 2 mm macro contacts at 8 x 8 orientation and sixty-four 1 mm micro contacts in between, providing 10 mm center-to-center spacing between macro contacts and 5 mm center-to-center spacing between micro/macro contacts, PMT corporation, Chanassen, MN). Recordings were made using one of two amplifier types: NicoletOne amplifier (Natus Neurologics, Middleton, WI), bandpass filtered from 0.16-250 Hz and digitized at 512 Hz. Neuroworks Quantum Amplifier (Natus Biomedical, Appleton, WI) recorded at 2048 Hz, bandpass filtered at 0.01682.67 Hz and then downsampled to 512 Hz. A two-contact subdural strip facing toward the skull near the craniotomy site was used as a reference for recording and a similar two-contact strip screwed to the skull was used for the instrument ground. Electrocorticography and experimental signals (trigger pulses that mark the appearance of visual stimuli on the screen, microphone signal from speech recordings and auditory playback signal that was heard by the patients through earphones) were acquired simultaneously by the EEG amplifier in order to provide a fully synchronized dataset. Recorded microphone and feedback signals were analyzed to ensure that the temporal delay manipulation by our MATLAB code produced the intended delay.

### Electrode localization

Electrode localization in subject space as well as MNI space was based on co-registering a preoperative (no electrodes) and postoperative (with electrodes) structural MRI (in some cases a postoperative CT was employed depending on clinical requirements) using a rigid-body transformation. Electrodes were then projected to the surface of cortex (preoperative segmented surface) to correct for edema induced shifts following previous procedures [27] (registration to MNI space was based on a non-linear DARTEL algorithm [28]). Within subject anatomical locations of electrodes was based on the automated FreeSurfer segmentation of the subject’s pre-operative MRI. All middle and caudal superior temporal gyrus electrodes were grouped as superior temporal gyrus (STG), all parsopercularis and pars triangularis electrodes were grouped as inferior frontal gyrus (IFG) electrodes. Precentral electrodes with a z coordinate smaller than ±40 were grouped as ventral precentral gyrus (vPreCG), and those with a z coordinate larger than or equal to ±40 were grouped as dorsal precentral gyrus (dPreCG) together with electrodes located in caudal middle frontal gyrus.

### Neural data analysis

A common average reference was calculated by subtracting the average signal across all electrodes from each individual electrode’s signal (after rejection of electrodes with artifacts caused by line noise, poor contact with cortex and high amplitude shifts). Continuous data streams from each channel were epoched into trials (from −1.5 s to 3.5 s for word stimuli and from −1.5 s to 5.5 s for sentence stimuli with respect to speech onset). Line noise at 60, 120 and 180 Hz were filtered out and the data was transformed to time-frequency space using the multitaper method (MATLAB, FieldTrip toolbox) with 3 Slepian tapers; frequency windows from 10 to 200 Hz; frequency steps of 5 Hz; time steps of 10 ms; temporal smoothing of 200 ms; frequency smoothing of ±10 Hz. The high gamma broadband response (70-150 Hz) at each time point following stimulus onset was measured as the percent signal change from baseline, with the baseline calculated over all trials in a time window from −500 to −100 ms before stimulus onset. High gamma response duration for each electrode was measured by calculating the time difference at full width quarter maximum of the response curve.

### Electrode selection

We recorded from a total of 1693 subdural and 608 depth electrode contacts in 15 subjects. Electrodes were examined for speech related activity defined as significant high gamma broadband responses. For auditory repetition and DAF word-reading tasks, electrodes that showed significant response increase (p < 10^-4^, unpaired t-test) either before (−0.5 to 0 s) or after speech onset (0 to 0.5 s) with respect to a baseline period (−1 to −0.6 s) and at the same time had a large signal-to-noise ratio (μ/σ > 0.7) during either of these time windows were selected. For DAF sentence-reading task, the same criteria were applied, except the time window after speech onset was longer (0 to 3 s). Electrode selection was first performed for each task separately, then electrodes that were commonly selected for both tasks were further analyzed.

### Clustering analysis

Non-negative matrix factorization (NMF) was used to identify major response patterns across different brain regions during speech production. NMF is an unsupervised dimensionality reduction (or clustering) technique that reveals the major patterns in the data without specifying any features [29]. We performed the clustering analysis using the data from the word-reading with DAF task, since this data set contained a large number of trials. We combined responses from all subjects by concatenating trials and electrodes forming a large data matrix A (electrodes-by-timepoints). Matrix A was factorized into two matrices W and H by minimizing the root mean square residual between A and W*H (nnmf function in MATLAB). Factorization was performed based on two clusters to represent the two major predicted speech related components in the brain; auditory and motor.

### Dynamic time warping analysis

For each trial of the DAF sentence-reading task, the speech spectrogram was averaged across frequencies. Then, the mean spectrograms were averaged across trials of the same sentence stimuli (e.g. averaged over Sentence #1 trials). Dynamic time warping (DTW) was performed separately for different sentence stimuli. The first 900 time points of the resulting DTW paths were applied to the neural response signal at each trial (representing approximately 9 seconds in the common warped time). Finally, the transformed neural responses were averaged across trials for each sentence stimuli. This procedure was performed to compare two conditions that resulted in the largest neural response difference (no delay versus 200 ms delay).

### Statistical analysis

The effect of DAF on speech behavior was determined by performing one-way ANOVA across subjects using articulation duration of words and sentences as the dependent variable and delay condition as the independent variable. To determine a significant difference in the amplitude of neural response between conditions, the average high gamma activity in a specified time window was compared by performing one-way ANOVA across all trials in all electrodes using delay condition as the independent variable. Similarly, a significant difference in the duration of neural response was determined by performing one-way ANOVA across subjects using response duration as the dependent variable and delay condition as the independent variable. Significance levels were computed at a p-value of 0.01. To compare sensitivity to DAF for word and sentence-reading, sensitivity indices were compared across electrodes using an unpaired t-test. To reveal how response enhancement to DAF changed across time during the sentence-reading task, we performed a one-way ANOVA at each time point using the neural response in each electrode as a dependent variable and delay as an independent variable. We performed a permutation test at each timepoint to assess a significance threshold for the F-value. We shuffled the delay condition labels 1000 times and performed an ANOVA for each timepoint at each iteration, then we set the threshold the 0.999 quantile of the F-value distribution.

## Abbreviations

DAF: Delayed auditory feedback
dPreCG: Dorsal precentral gyrus
DTW: Dynamic time warping
ECoG: Electrocorticography
IFG: Inferior frontal gyrus
NMF: Non-negative matrix factorization
PostCG: Postcentral gyrus
SMG: Supramarginal gyrus
STG: Superior temporal gyrus
vPreCG: Ventral precentral gyrus

## Acknowledgement

We thank Zhuoran Huang and Qingyang Zhu for their assistance in analyzing voice recordings of the subjects.

## Notes

### Competing Interest Statement

The authors have declared no competing interest.

